# Ovipin: a new antimicrobial peptide from chicken eggs *Gallus gallus*

**DOI:** 10.1101/2021.09.28.462162

**Authors:** Sandra Regina dos Santos, Antonio Miranda, Pedro Ismael da Silva Junior

## Abstract

The intensive and indiscriminate use of antibiotic has increased cases of microorganisms resistance and becoming a worldwide public health problem. In the last years, from natural sources such as hen’s egg, have received special attention in the discovery of new bioactive compounds. This study aims to identify and characterize a new peptide from chicken egg of *Gallus gallus domesticus*. The chicken egg was subjected the extraction acid, the supernatant was prefractionated in Sep-Pak column and fractionated peptide by reverse-phase high-performance liquid chromatography (RP-HPLC). The antimicrobial activity of the fractions were evaluated through liquid growth inhibition assays. The molecular weight and amino acid sequence was determined by mass spectrometry (MS/MS), the characaterization performed by bioinformatics analysis with Peaks® tool and comparision with the NCBI and UniProt-SwisProt database. The physicochemical parameters of the samples were evaluated with online programs. One fraction named Ovipin peptide, showed antimicrobial activity against Gram-positive bacteria (*Micrococcus luteus* – MIC 1.94 µM) and Filamentous fungus (*Aspergillus niger* - MIC 31.01 µM). The minimum inhibitory concentration against *Cryptococcus neoformans* VNI (WM 148) Yeast was 15.51 µM, this microorganism an opportunistic yeast and mainly binds in immunosuppressed or immunocompromissed people. Ovipin is a hydrophoic peptide and not cause significant hemolytic effect against human erythrocytes. Ovipin primary sequence is YVSPVAIVKGLNIPL and a molecular weight of 1,581.94 Da. Ovipin shows 100% and 93.3%, respectively, sequence similarity with the fragments Apolipoprotein B of *Gallus gallus* and Apolipoprotein B of five others species of Aves. Our data suggest that Ovipin peptide could be a potential therapeutic candidate to be used an alternative against infections developed by resistant microorganisms, mainly in the fight against *Cryptococcus neoformans* opportunistic yeast.

## 1 Introduction

Antimicrobial drugs are compounds discovery in differents sources and utilized in medicine to control the action of bacteria, fungi, viruses and parasites. However, these microorganisms naturally developed mechanisms to overcome the efects of antimicrobial drugs and ensure your survival (O’Neill, 2016), the antimicrobial resistance (AMR) is a serious global public health and socialeconomic problem because limit the options treatment against infections disease (WHO, 2022; Ben-Ami, Kontoyiannis, 2021). Selection pressure, excessive use of antibiotics in food (agriculture), animal production and humans, poor sanitation/hygiene, are some reasons which may favor the increase of antimicrobial resistance (Aslam *et al*., 2018). Data estimated that in 2019 occurred 4.95 million deaths associated with AMR and 1.27 million of the deaths related to bacterial resistance, it has been demonstrated that the main resistant bacteria are *Eschericchia coli, Staphylococcus aureus, Klebisiella pneumoniae, Streptococcus pneumoniae, Acinetobacter baumanii* and *Pseudomonas aeruginosa* (Muray *et al*., 2022).

How antimicrobial resistance is a global concern for the medical community and global leaders (Laxminarayan *et al*., 2016), the antimicrobial peptides (AMPs), are bioactive molecules with small molecular weight (<100 amino acid residues), and they are the first barrier of defense against invading pathogenic microorganisms (Li *et al*., 2022), several studies have been carried out focusing on these molecules as an alternative to conventional antibiotics. The AMPs have broad spectrum antimicrobial and low natural resistance, it can be obtained from different sources, like as plants, animal, microorganisms and foods (Liu *et al*., 2021; Abdel-Razek *et al*., 2020). Doderlin peptide was isolated and charactherized from *Lactobacillus acidophilus* bacteria, this peptide was effective against yeast *Candida albicans* strains (Silva *et al*., 2023). In the mucus *Limacus flavus* slug were isolated and identified two antimicrobial peptides with antimicrobial activity against Gram-positive bacteria (Hayashida, Silva Junior, 2021).

Antimicrobial peptides act through the electrostatic interaction of their positive charges present in the plasma membrane of microorganisms, this interaction triggers changes in membrane permeability followed by formation pore with intracellular leakage and death of the microorganims (Mba, Nweze, 2022). Some peptides act on intracellular targets (Li *et al*., 2022), such as Sarconesin II (Díaz-Roa *et al*., 2019) and Rondonin (Riciluca *et al*., 2021), suggest these peptides act on nucleic acids of *Eschericchia coli* and *Candida albicans*, respectivelly.

Foods are also sources for identifying bioactive compounds (Liu *et al*., 2021). In the work of Torres *et al*., (2021) was isolated and identified two antimicrobial compouns from garlic that were effective against *Streptococcus agalactiae* strains. The chicken eggs is a complete food because it has a good nutritional balance, such as water, proteins, lipidis, vitamins and minerals, and it a source of biologically active compounds (Réhault-Godbert, Guyot, Nys, 2019).

Different types of proteins present in egg white or yolk were described in the literature demonstrated antimicrobial action, of lysozyme, ovotransferrin, avidin, roboflavin-binding protein, cystatin, ovostatin, ovomucin, phosvitin, Tenp and ovocalyxin-36 (Nys, Bain, Van Immerseel, 2011). Egg-derived peptides obtained these proteins have demonstrated antibacterial and antifungal activity against microorganism resistants (Réhault-Godbert, Guyot, Nys, 2019), like as Mine et al., (2004) identified antimicrobial peptides of enzymatic hydrolisis of lysozyme; OVTp12 peptide from hydrolysate ovotransferrin was effecitve against Gram-positive and Gram-negative bactéria (Ma *et al*., 2020) and Pimchan *et al*., (2023) isolated, identified, evaluted antibacterial peptide from egg yolk hydrolysate.

So, the aim of this work is to performed the bioprospecting on chicken eggs *Gallus gallus domesticus*, with focus in the isolation of new antimcrobial peptides by liquid chromatography, bioassay to determine antimicrobial activity potential, and characterization by mass spectrometry and bioinformatics analysis.

## 2 Materials and Methods

### 2.1 Acid and Solid-Phase Extraction

In this study chicken eggs without the shell of the species *Gallus gallus domesticus* were utilized. They were purchased from the commercial market network in the city of São Paulo/SP.

The total chicken eggs crude were submitted to acid extraction in the presence of Acetic Acid 2 mol.L^−1^, with constant agitation for 60 min at 4° C. After the stirring time, the insoluble material was separated by centrifugation at 3113 x g for 30 min and the supernatant obtained was injected into coupled Sep-Pack_18_ cartridges (Waters Associates – 20 cc vac cartridge, 5 g, and particle weight 55- 105 µm) equilibrated in trifluoroacetic acid (0,1% TFA). The supernatant was pre-fractionated in three different acetonitrile (ACN) concentrations (5%, 40%, and 80% of ACN), using vacuum manifold system equipment (SPE 12 positions aho 6023 - Phenomenex) and then were lyophilized.

### 2.2 Reverse-Phase High-Performance Liquid Chromatography (RP-HPLC)

The fractions, 5%, 40% and 80% of ACN with 2 g each, were reconstituted in 5 mL 0.1% TFA and subjected by reverse-phase high-performance liquid chromatography (RP-HPLC) was carried out at room temperature on a Shimadzu LC-8A system. The column was a Shim-Pack Preparative ODS C-18 (20 mm x 250 mm; particle weight 10 µm; pore weight 300 Å). The elution was performed with a linear ACN gradient, equilibrated in 0.1% TFA for 60 min at a flow rate of 8.0 mL/min (0 to 20%) for the fraction eluted in 5%, 2 to 60% for the fraction in 40%, and 20 to 80% fraction eluted in 80%. The ultraviolet (UV) absorbance of the effluent was monitored at 225 nm. The fractions were manually collected, were vacuum-dried and suspended in 1 mL in ultrapure water for antimicrobial activity assays.

The molecule described in this work, fraction 21, was subjected to a second step of fractionation using linear gradient of ACN (39 to 49%) with 0,05% TFA at a flow rate of 2 mL/min for 60 min on the same column Shim-Pack Preparative ODS C-18 (20 mm x 250 mm; particle weight 10 µm; pore weight 300 Å). The fractions were hand-collected, were vacuum-dried, suspended in 500 µL in ultrapure water and it was quantified based on absorbance 205 nm using a NanoDrop 2000 spectrophotometer (Thermo Fischer Scientific, Waltham, MA, USA).

### 2.3 Microorganisms

Fungal and bacterial strains were obtained and stored in the freezer at -80°C in the Laboratory for Applied Toxinology (LETA), of the Butantan Institute (São Paulo, Brazil). *Staphylococcus aureus* ATCC 29213 (Gram-positive bacteria), *Escherichia coli* D31 (Gram–negative bacteria); *Aspergilus niger* (Filamentous fungus – isolated from bread); and *Cryptococcus neoformans* yeast strain A serotype (CFP registration 55 (WM 148 = VNI) were used. The *Cryptococcus neoformans* strain was granted by Dra. Marcia Melhem from the Adolfo Lutz Institute, Brazil - São Paulo.

### 2.4 Antimicrobial Assays and Minimal Inhibitory Concentration (MIC)

Antimicrobial activities of the fractions were evaluated by a liquid growth inhibition assay (Bulet *et al*., 1993). A suspension of the microorganisms collected in the mid-logarithmic growth phase was used. Bacteria strains were cultured in poor broth nutrient medium (PB: 1.0 g peptone in 100 mL of water containing 86 mM NaCl at pH 7.4; 217 mOsM); for the fungi and yeast strains were cultured in potato dextrose broth (1/2-strength PDB) (1.2 g potato dextrose in 100 mL of H_2_O at pH 5.0; 79 mOsM). Determination of antimicrobial activity was performed using 5-fold microtiter broth dilution assay in 96-well sterile plates at a final volume of 100 µL. A mid-log phase culture were diluted to a final concentration of 5 x 10^4^ CFU/mL for bacteria and 5 x 10^5^ CFU/mL for fungus and yeast (Bulet, 2008; Hetru and Bulet, 1997). Sterile water and PB or PDB were used as growth control, and Streptomycin antibiotic (10.0 mg/mL) was used as growth inhibition control. Microtiter plate were incubated for 18 and 24 h for bacteria and fungi/yeast, respectively, at 30°C; growth inhibition was determined by measuring absorbance at 595 nm. The minimum inhibitory concentration (MIC) are expressed as the interval of the concentration [a]–[b], where [a] is the highest tested concentration at which the microorganism grow, and [b] is the lowest concentration which causes a growth inhibition of 100% (Bulet *et al*., 1993). The assay was performed using a serial dilution and duplicate in 96-well sterile plate; at 20 µL stock solution was used in each microtiter plate well and added to 80 µL of the microorganism dilution (Silva *et al*., 2000; Lorenzini *et al*., 2003; Riciluca *et al*., 2012).

### 2.5 Determination of Hemolytic Activity

The hemolytic activity of the fractions with antimicrobial activity was assessed against human erythrocytes from a health adult donor. The Ethics Committee was the University of São Paulo School of Medicine (USP - FMUSP), n° 18794419.7.0000.0065. The erythrocytes were collected in 0.15 M citrate buffer, pH 7.4, and washed three times by centrifugation (700 x g, 10 min, 4°C) with 0.15 M phosphate buffer saline (PBS – 35 mM phosphate buffer, 0.15 M NaCl, pH 7.4), and ressuspended in PBS to a final 3% (v/v) concentration.

Aliquots of 50 µL of each fraction were added to 50 µL of 3% (v/v) suspension erythrocytes in the well of U – shaped bottom plates and incubated for 1h at 37° C (with a final volume of 100 µL). The supernatant was first collected and hemolysis was measuring the absorbance at 405 nm of each well in a Victor^3^ (1420 Multilabel Counter/ Victor^3^, Perkin Elmer). The hemolysis percentage was expressed in relation to a 100% lysis control (erythrocytes incubated with 0.1% triton X-100) (Sigma-Aldrich, St. Louis, MO, USA); PBS was used as negative control. The fractions was tested serial dilution and duplicate, and calculation was made according to the fallowing equation: %hemolysis = (Abs sample – Abs negative)/(Abs positive – Abs negative) (Riciluca et al., 2012; Díaz-Roa et al., 2018; Segura and Silva Junior, 2018).

### 2.6 Mass Spectrometry Analysis

The fractions that showed antimicrobial activity were analyzed by positive mode mass spectrometry LC-MS/MS on a LTQ XL (Thermo Scientific). The fractions (10 µL of which) were concentrated and ressuspended in 10 µL Formic Acid 0,1%. The liquid chromatography was perfomed in a C18 column (13 cm; 75 µm, i. d.x 100 mm – 3 µm from Waters), using a 5 to 95%B in 25 minutes with solvents A: 15% in acidified water (0.1% TFA) and B: 95% in acidified water (0.1% TFA) for 2 min; with flow of 300 *n*L/min, voltage of 2.5 kV at 200°C. The full scan was performed with m/z of 100 a 2000, occurring MS/MS–CID, and parameters of five most intense ions, with isolation width of 2 m/z at CID (collision energy at 35).

### 2.7 Database Search

The resulting spectra in “*.RAW*”, format were collected and processed using MSconvert software (Chambers et al., 2012). Thus, they were converted to the mascot generic format (“*.MGF*”) and thus analyzed into database searches using the Mascot tool (Perkins et al., 1999). The MS/MS peak list files were submitted to an in – house version of the Mascot server (Matrix Science, United States) and screened against the Uniprot database. PEAKS Studio Software (version 8.5, Bioinformatics Solutions Inc., Waterloo, Ontario, Canada) *de novo* sequencing/database search used for establishing sequence. Analysis involved 10 ppm error tolerance for precursor ions and 0.6 Da for fragment ions. Oxidation was considered a variable modification.

The physicochemical parameters of which the fraction (such as the total number of positively and negatively charged residues, molecular weight, and theoretical pI) were calculated using the ProtParam tool, available through the portal ExPASy of the Swiss Institute of Bioinformatics website (https://web.expasy.org/protparam/) (Gasteiger et al., 2005).

## 3 Results

### 3.1 Chicken Eggs Fractionation and Antimicrobial Activity

The crude extract from chicken eggs was pre-fractionated in column Sep-Pak Cartridge, obtained three elutions 5%, 40% and 80% ACN. By RP-HPLC, samples eluted 5% and 40% ACN concentration not show antimicrobial activity and only six fractions (9, 11, 20, 21, 22 and 24) eluted 80% ACN concentration (**Figure 1A**), showed antimicrobial activity in liquid growth antimicrobial assay (**Table 1**). The fractions 20, 21 and 22 were effectiveness against all tested microorganisms.

**Table 1.** Antimicrobial activity of six fractions from Chicken eggs.

| Fractions | Microorganisms |  |  |  |
| --- | --- | --- | --- | --- |
|  | Fungus |  | Bacteria |  |
|  | Filamentous<br>*A. niger* | Yeast<br>*C. neoformans* | Gram-positive<br>*S. aureus* | Gram-negative<br>*E. coli* |
| 9 | - | + | - | - |
| 11 | + | + | - | - |
| 20 | + | + | + | + |
| 21 | + | + | + | + |
| 22 | + | + | + | + |
| 24 | + | + | - | - |

**FIGURE 1:**
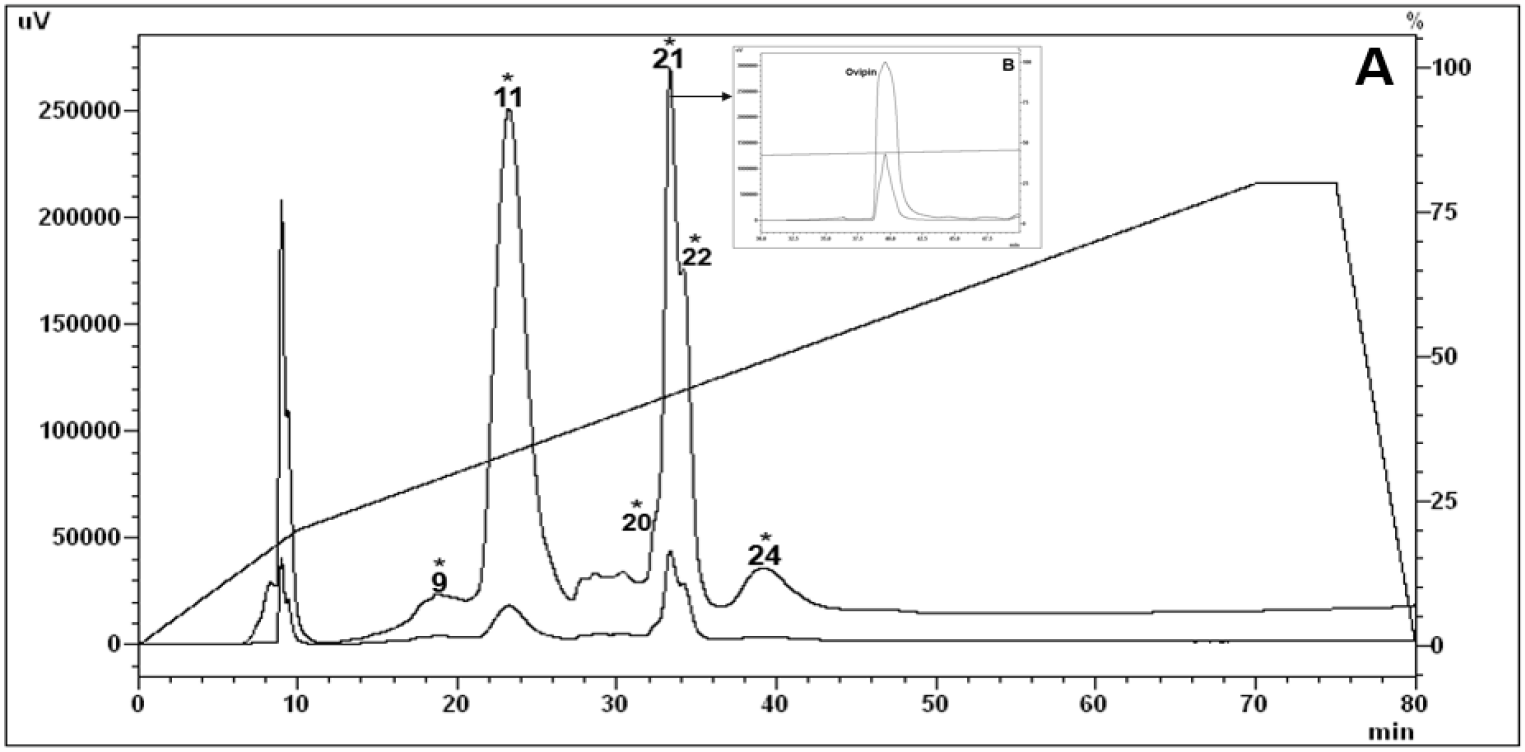
Reversed-phase high-performance liquid chromatograph (RP-HPLC) of the Chicken eggs *Gallus gallus domesticus* extract eluted with 80% acetonitrile (ACN). After the pre-fractionation in a Sep-Pak C18 column, the fraction was eluted with 80% ACN were subjected to a Shim-Pack ODS C18 (20 x 250 mm, 300 Å, 10 µm); Solvents System: Solvent A: 0.1% TFA/H_2_O and Solvent B: 0.1%TFA/ACN with a linear gradient 20% to 80% ACN, 60 min at 8 mL/min flow rate. (**A**)The fraction 21 indicated by an asterisk (*) had antimicrobial activity against microorganisms tested. (**B**) Ovipin, indicated with an arrow, using same column and solvents system: Solvent A: 0.05% TFA/H_2_O and Solvent B: 0.05%TFA/ACN. Ovipin was eluted from linear gradient 39% to 49% ACN, for 60 min at 2 mL/min flow rate. Both runs the absorbance was monitored at 225 nm.

### 3.2 Minimum Inhibitory Concentration (MIC) of Ovipin peptide

The fraction 21 eluted with retention time range of 32.7-34.1 min (**Figure 1A**), it was submitted to a second step of fractionation and showed only one a pronounced antimicrobial fraction: named Ovipin (**Figure 1B**). The name to refer the chicken eggs species *Galus gallus domesticus*. The initial concentration of the native Ovipin peptide applied in the serial dilution was of 124,05 µM (182 µg/mL), and MIC value of Ovipin peptide against *M. luteus* A270 1,94 µM (2,84 µg/mL), *A. niger* 31,01 µM (45,5 µg/mL), *C. neoformans* VNI (WM 148) 15,51 µM (22,75 µg/mL). And the *E. coli* D31 was the least sensitive to this peptide (**Table 2**).

**Table 2.**
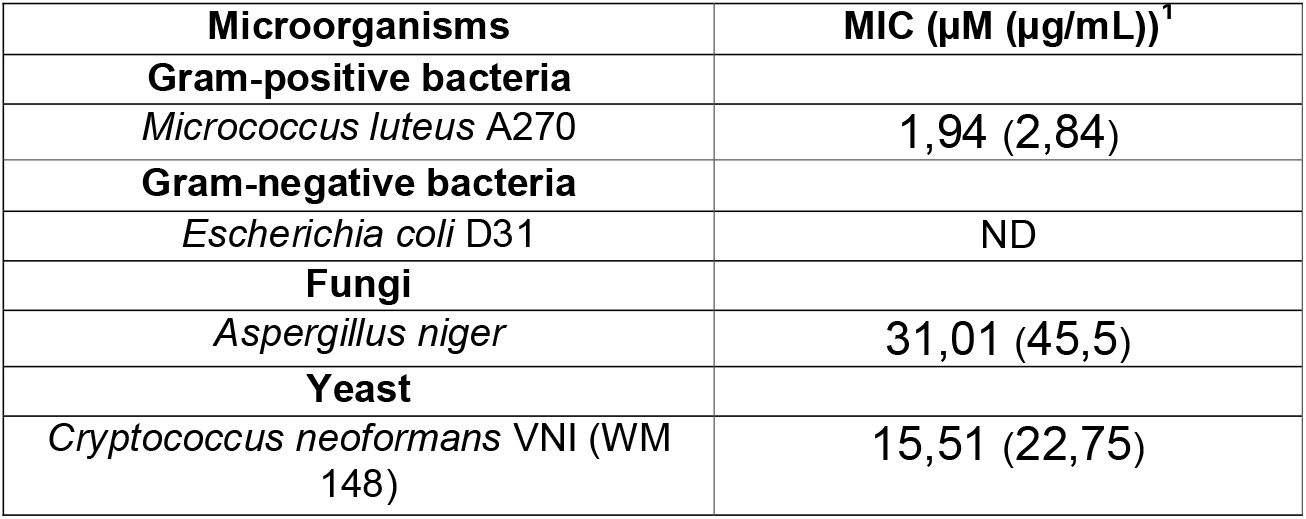
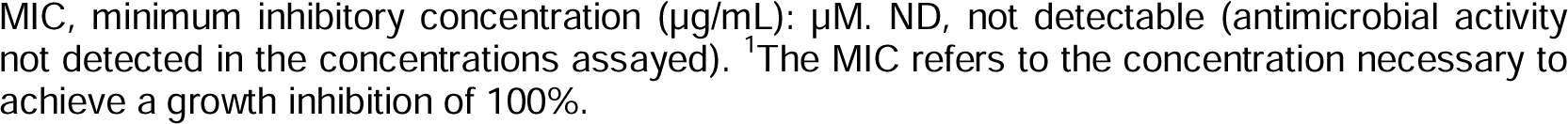
Antimicrobial activity spectrum of Ovipin peptide.

| Microorganisms | MIC (µM (µg/mL))<sup>1</sup> |
| --- | --- |
| **Gram-positive bacteria** |  |
| *Micrococcus luteus* A270 | 1,94 (2,84) |
| **Gram-negative bacteria** |  |
| *Escherichia coli* D31 | ND |
| **Fungi** |  |
| *Aspergillus niger* | 31,01 (45,5) |
| **Yeast** |  |
| *Cryptococcus neoformans* VNI (WM 148) | 15,51 (22,75) |

MIC, minimum inhibitory concentration ( $\mu\text{g/mL}$ ): $\mu\text{M}$ . ND, not detectable (antimicrobial activity not detected in the concentrations assayed). <sup>1</sup>The MIC refers to the concentration necessary to achieve a growth inhibition of 100%.

### 3.3 Hemolytic Activity

To determine the effect of the Ovipin peptide on human erythrocytes at the antimicrobial concentrations, its hemolytic activity was tested. After incubating red blood cells from a healthy donor with Ovipin (124.05 µM up to 0.24 µM) concentrations for 1 h at 37 °C. No hemolytic activity was observed, demonstrated that Ovipin does not cause significant hemolytic effect against of human erythrocytes within these concentrations (**Figure 2**) (Yacoub *et al*., 2014).

**FIGURE 2:**
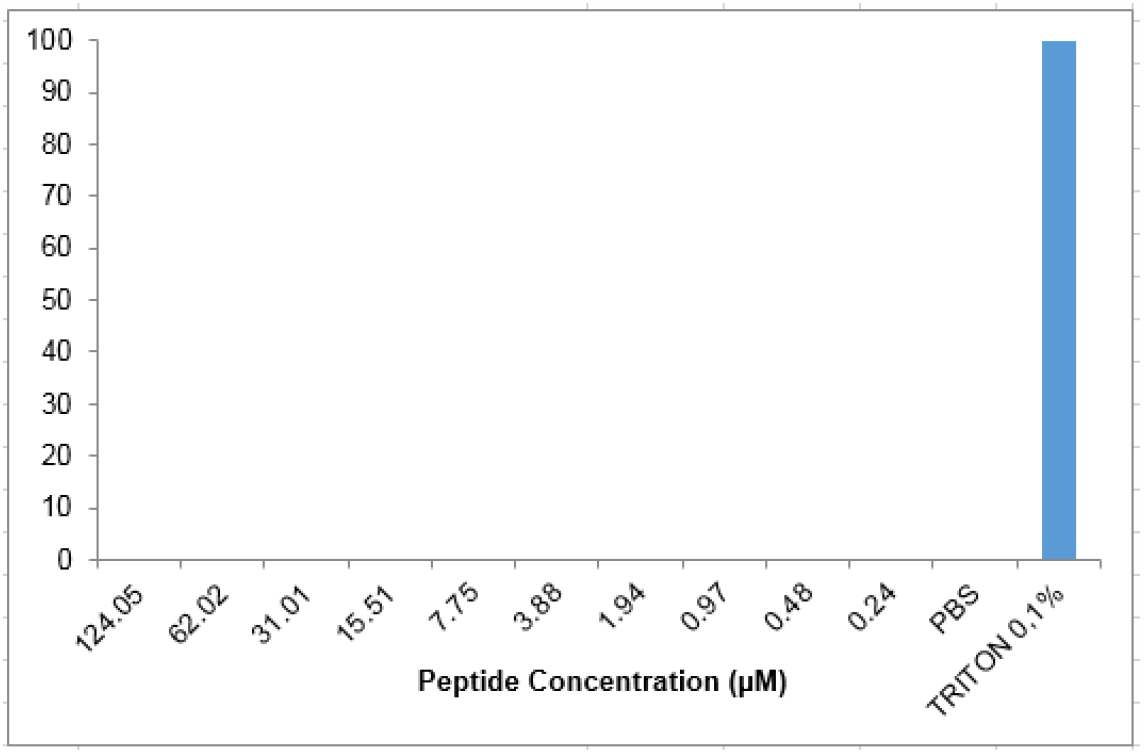
Hemolytic Activity of the Ovipin peptide incubated with human erythrocytes in different concentrations of the peptides. Serial dilution and duplicated was performed, for 1 h at 37°C. The hemolysis percentage was expressedin relation to 0% hemolysis with PBS and 100% lysis control (0.1% triton X-100)

### 3.4 Mass Spectrometry and Characterization of Ovipin Peptide

Mass spectrometry of the Ovipin peptide was analyzed using the in-house MASCOT® software. This tool revealed the amino acid sequence YVSPVAIVKGLNIPL, molecular weight of 1,581.9494 Da, and using the Pepdraw tool revealed the primay structure (**Figure 3A, 3B 3C**). By the deconvolution of the peptide and analysis of the data in Peaks DB software, the PEAKS *de novo* sequencing showed, as well as the Mascot tool, 15 amino acid sequence do not having post-translational modification (PTM). Collision-induced dissociation (CID) spectrum from mass/charge (m/z) of its double charged ion give [M + 2H]^2 +^, m/z 792.3484 (**Figure 4**).The Peaks DB tool revealed which primary sequence of the Ovipin derived from Apolipoprotein B (ApoB) and cover 2% of the whole protein sequence (**Figure 5**).

**Figure 3:**
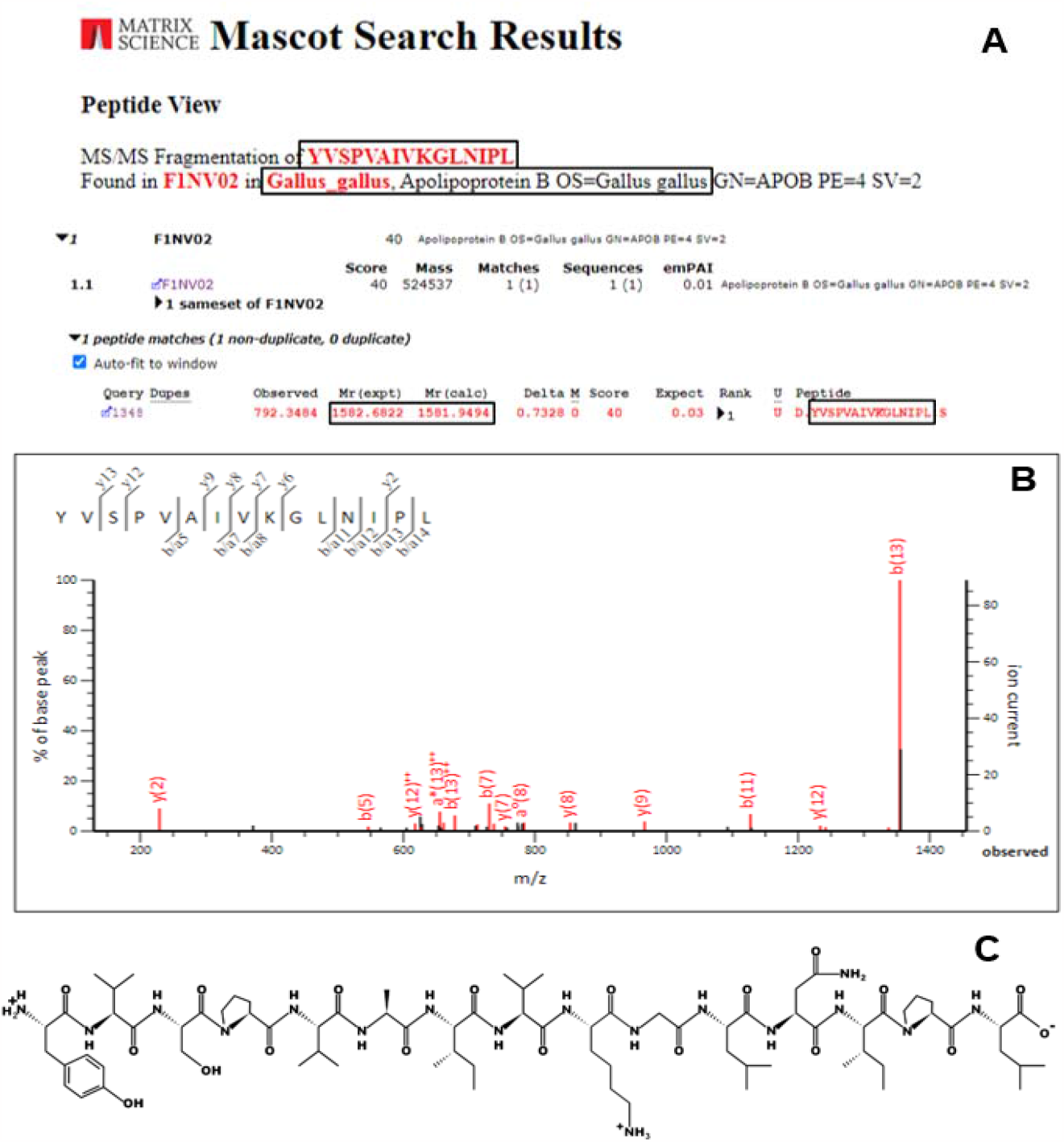
Result obtained in searches databases using the tool Mascot**®** (Matrix Science) and PepDraw tool. (**A**). The Ovipin peptide is similar with a fragment from the *Gallus gallus* Apolipoprotein B (APO B). (**B**) MS/MS spectra generated from CID fragementation of Ovipin and submitted to Mascot search, revealed the “YVSPVAIVKGLNIPL” sequence generated by deconvolution of “b” and “y” peptide ions. (**C**) The primary structure of the Ovipin peptide obtained by PepDraw tool.

**Figure 4:**
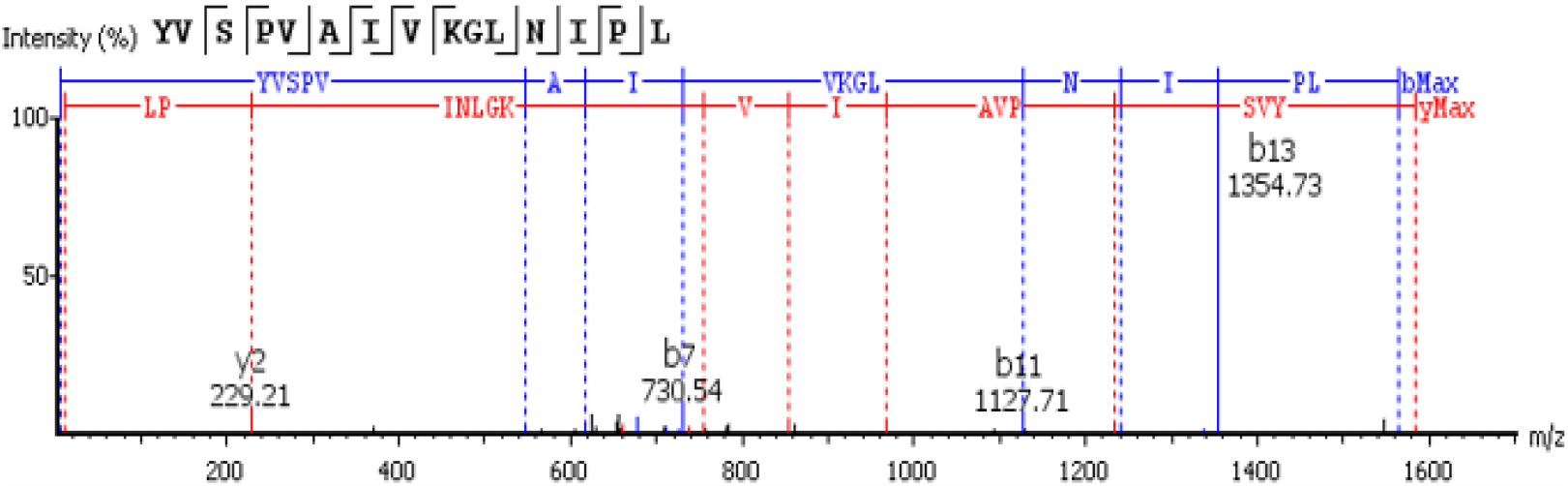
Collision-induced dissociation (CID) spectrum of *de novo* sequencing antimicrobial peptide, Ovipin. The ions belonging to –y (red) and –b (blue) series indicated in the spectrum correspond to the amino acid sequence of the peptide: YVSPVAIVKGLNIPL. The fragments of the sequenced peptide are represented by standard amino acid code letters.

**Figure 5:**
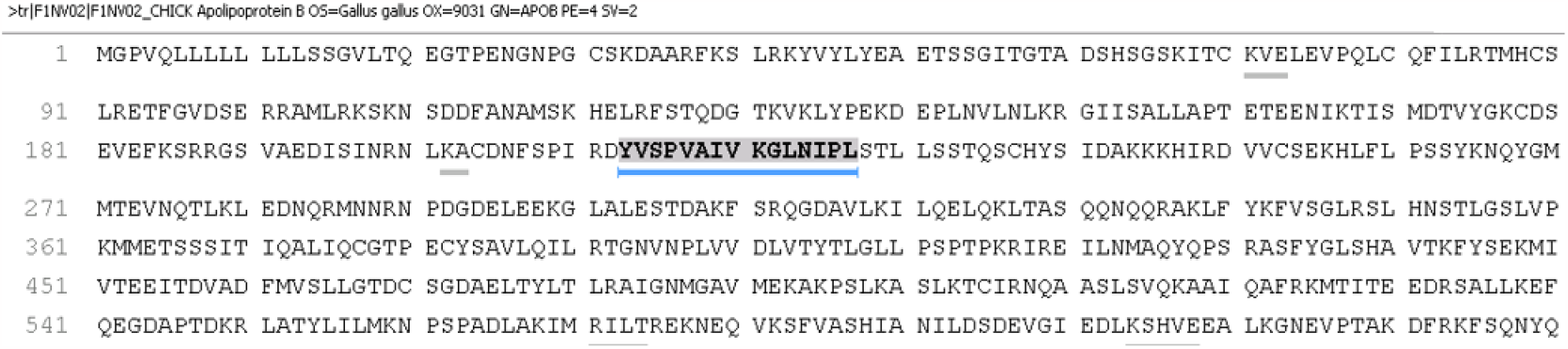
Peptide spectrum match indicated by a blue line below the sequence of the Apolipoprotein B of *Gallus gallus*. Ovipin covered 2% of the whole protein sequence.

The amino acid sequence Ovipin was analyzed in the APD database (**Figure 6**), it identified highest similarity (47,37%) with the temporin-1SKa from the Asia frog *Rana sakuaii*, temporin-1GY (43,75%) from the Asia frog *Pelophylax nigromaculatus*, peptide AN5-1 (41,18%) from *Paenibacillus alvei* AN5 and odorranain-R1 (40%), from frog *Odorrana gharami*. The Clustal Omega program was used to identify a multiple alignment analysis of the peptide sequence Ovipin, the searches showed 100% sequence alignments with the fragment Apolipoprotein B of *Gallus gallus*, and 93.3% of similarity with fragments Apolipoprotein B with others five species of Aves, being *Patagioenas fasciata monilis, Amazona aestiva, Tinamus guttatus, Aptenodytes forsteri* and *Egretta garzetta* (Figure 7).

**Figure 6.**
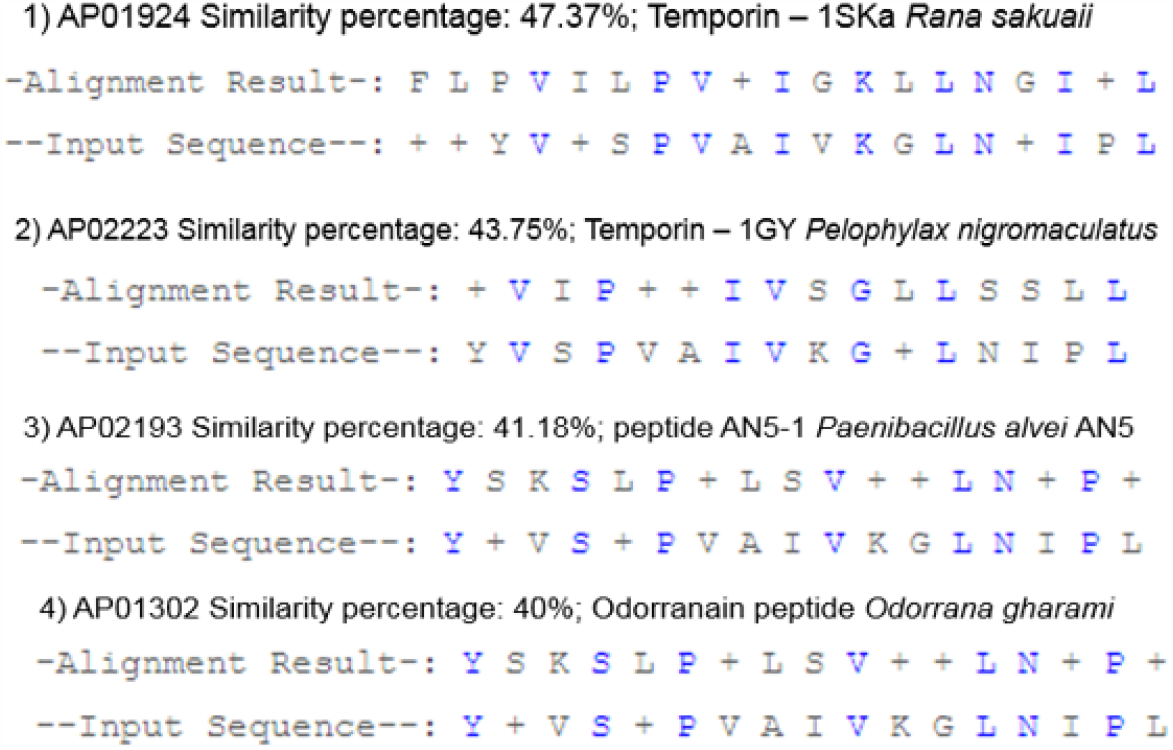
Porcentage alignments results of the Ovipin peptide with Antimicrobial Peptide Database (APD) (**Wang et al**., **2009**).

**Figure 7:**
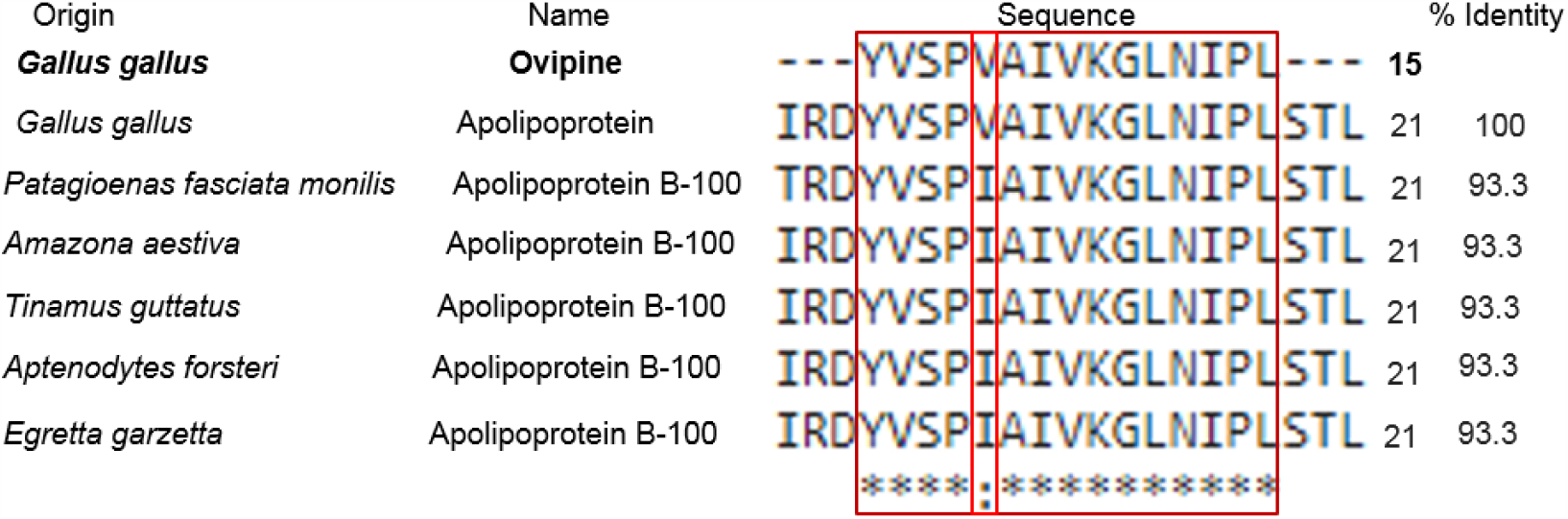
Alignment of the amino acid sequence of Ovipin. Using the Clustal Omega alignment, the Ovipin sequence was aligned with fragments of the Apolipoproteins of the *Gallus gallus* and others Aves species. Regions that show identical amino acid among all species are identified in dark red or (**\***), and minor modifications are show in light red (:). Clustal Omega (http://www.ebi.ac.uk/Tools/msa/clustalo/; accessed on 24 January 2020) modified manually.

### Physochemical Characteristics

The ExPASY’s (SIB Bioinformatics Resource Portal), PepDraw and Pep-Calc tools were used predicting Ovipin significant physicochemical characteristics (**Table 3**). The sequence Ovipin contains 15 amino acids (aa), were predicted to be hydrophobic aa residues as two Ile (I), two Leu (L), three Val (V), one Ala (A), with total positively charge (+1) and basic nature (isoelectric point 9.54). The instability index (II) of Ovipin was of the 65.60, classified this peptide as may be unstable (instability is than greater 40).

**Table 3:** Ovipin theoretical physicochemical properties.

| Ovipin peptide properties |  |
| --- | --- |
| Sequence: | YVSPVAIVKGLNIPL |
| Lenght: | 15 |
| Mass: | 1581.9494 |
| Isoelectric point (pI): | 9.49 |
| Net charge at pH: | 1 |
| Hydrophobicity: | + 7.11 Kcal*mol<sup>-1</sup> |
| Extinction coefficient: | 1280 M<sup>-1</sup> cm<sup>-1</sup> |
| Total hydrophobic ratio | 53% |
| Aliphatic Index (AI) | 168,60 |
| GRAVY | 1,193 |
| Estimated solubility: | Poor water solubility |

Physicochemical parameters were calculated using ExPASy PepDraw and Pep-Calc.com

The aliphatic index (AI) is a parameter important used for avalied thermostability of protein according the relative volume occupied by aliphatic side chain, aliphatic amino acids are also hidrophobic nature (IKAI, 1980). The aliphatic index of Ovipin showed 168.60, indicated this peptide is thermally stable because it contain hidrophobic amino acids in your structure (Panda, Chandra, 2012). The grand average of hidropathicity (GRAVY) plays role in calculate hidropathy value of the amino acid sequence and evaluate the solubility of the proteins, indicating hidrodrophibic (positive GRAVY) and hidrophilic (negative GRAVY) (Silva *et al*., 2023; Kyte, Doolittle, 1982). Ovipin showed hydrophobic ratio of 53%, positive GRAVY with value of 1,193 indicated be hydrophobic and suggesting poor water solubility. ExPASy’s ProtParam tool predicted that the peptide would intact for up 2.8 hours in mammalian reticulocytes (*in vitro*), 10 min in yeast (*in vivo*) and 2 min in *Escherichia coli* (*in vivo*).

## 4 Discussion

People immunosupressed or immunocompromised system are those most affected by microorganisms resistant to conventional antibiotics (Vitiello *et al*., 2023), increasing the spectrum of untreatable infections and generated a great worldwide public health problem (Gaglione *et al*., 2011; Blair *et al*., 2015). The accelerated growth of resistance to antimicrobials, new strategies and discovery of new efficient compounds mainly with antimicrobial action and that have low toxicity for human cells are urgently needed, to be used as tools for antimicrobial therapies (Vitiello *et al*., 2023; Díaz-Roa *et al*., 2018; Chernysh *et al*., 2015).

Chicken eggs are an easy to obtain natural source for research and identification of molecules with different functions. In the present work we are isolated by RP-HPLC six fractions which were capable inhibity the microorganisms tested. Ovipin peptide was inhibited the growth of *Micrococcus luteus* A270 (MIC = 1,94 µM (2,84 µg/mL)), *Aspergillus niger* (MIC = 31,01 µM (45,5 µg/mL)) and this peptide not was able inhibited the growth of *E. coli. M. luteus* in general it is founded in skin, mucous membranes and oropharynx, but in immunosuppressed patients this microorganism can triggered bacteremia infection (Martín Guerra *et al*., 2019). Invasive aspergillosis is also problem serious for a medicine because affect mainly immunocomppromissed individuals or patients which receive immunosuppressive treatments, as also solid organ transplantation (Sugui *et al*., 2014). In the study Yacoub *et al*., (2016), the synthetic chicken β-defensin-9 (sAvBD-9) was active against Gram-positive, Gram-negative bactéria and yeast, like also inhibited growth the *Aspergillus flavus* (MIC 3.2 µg/mL and MFC 3.2 µg/mL) and *Aspergillus niger* (MIC 3.2 µg/mL and MFC 6.25 µg/mL).

In the work Candido-Ferreira *et al*. (2017) was identified and characterized the Oligoventin peptide (8 amino acid residues and MW 1,061.4 Da) from the eggs of the Brazilian armed spider *Phoneutria nigriventer* (Ctenidae, Araneomorphae). Oligoventin showed antimicrobial activity (MIC range at 47 to 188.9 µM) against Gram-positive and Gram-negative bacteria, as also yeast (*Candida albicans* MDM8). Therefore, it is worth pointing out that both Ovipin and Oligoventin, although islolated from eggs of different animals, have antimicrobial action against opportunistic pathogens of great medical importance.

The antimicrobial activity observed of Ovipin (MIC = 15,51 µM (22,75 µg/mL)) was against *Cryptococcus neoformans* is very relevant because it an opportunistic pathogen which cause human infection such as cryptococcal meningits, with highly incidence of morbidity and mortality in immunocompromised individuals, as HIV/AIDS patients, cancer chemoterapy, hematologic and solid organ transplantation (Bermas, Geddes-McAlister, 2020; Roemer, Krysan, 2014). As well as Ovipin, Doderlin peptide from natural source *Lactobacillus acidophilus* extracts, was also effective against *C. neoformans* strains (MIC = 116-464 µg/mL (50-200 µM)) and inhibited the growth of *Candida* strains (SILVA *et al*., 2023). In general, molecules that have already been isolated and identified in chicken eggs showed antimicrobial action against bacterial and some fungal strains, but are still little explored the compounds identified in chicken eggs with activity against strains of *Cryptococcus spp*. And as shown in the priority list for pathogenic fungi with the greatest threat drawn up by the World Health Organization, *Cryptococcus neoformans, Candida auris, Candida albicans* and *Aspergillus fumigatus*, are the four microorganisms included in this list as critical priority, because they can cause invasive acute and subacute systemic fungal infections (WHO, 2022).There is great concern for cases of resistance to conventional antifungal drugs, polyenes, azoles and echinocandins, the Ovipin peptide may be a future alternative in the treatment of antifungal therapy, mainly against *Cryptococcus spp*. (Roemer, Krysan, 2014).

Through searches carried out in online databases, Ovipin demonstrated a high percentage of similarity with bacteria and frog peptides. It has identified highest similarity (47,37%) with the temporin-1SKa peptide extracted of the skin frog from *Rana sakuaii* (Suzuki *et al*., 2007), temporin-1GY (43,75%) of the skin frog from *Pelophylax nigromaculatus* (Song et al., 2013), peptide AN5-1 (41,18%) from *Paenibacillus alvei* AN5 (Alkotaini et al., 2013) and odorranain-R1 (40%), from frog *Odorrana gharami* (Li *et al*., 2007). Amphibians, like chicken eggs, are frequently exposed to many pathogens and studies have shown several isolated and identified antimicrobial peptides or pharmacological peptides from frogs with diverse function, mainly against multi-resistant microorganisms (Hayashida, Silva Junior, 2021; Xiao *et al*., 2011).

The results of the alignments using Clustal Omega, indicated the Ovipin sequence to have high similarity, 100% and 93.3%, with Apolipoprotein B-100 fragments from *Gallus gallus* and five species of Aves, like as *Patagioenas fasciata monilis, Amazona aestiva, Tinamus guttatus, Aptenodytes forsteri* and *Egretta garzetta*. Apolipoproteins are recognized as source of bioactive peptides showed broad antimicrobial or antiviral activity, the Apoliproteins B is formed by two low density lipoprotein (LDL) of name region A and region B. Region B is more conserved between species (Kelly et al., 2010). In hen egg yolk, the molecular weight of Apo-B protein is 500 kDa and 64% of the similarity with human apolipoprotein B-100 precursor (Jolivet *et al*., 2006).

Peptides derived from egg yolk hiydrolysate showed antibacterial action against *Staphylococcus aureus* ATCC 29213 and *Salmonella typhimurium* TISTR 292 at MIC values of 0.5 to 1 mmol/L. These peptides demonstrated cationic and anionic nature, and showed 100% similarity to fragments with Apolipoprotein B in egg yolk of the *Gallus gallus* (Pimchan et al., 2023), samilar result of Ovipin peptide. The study Zambrovicz *et al*., (2015) was identified four peptides with molecular weight ranging from 1,210.62 to 1,677.88 Da obtained from Apoliproteins B, Vitellogenin-2 and Apovitelenin-1, all proteins localized in chicken egg yolk. In relation the peptides identified from Apolipoprotein B, one peptide showed amino acid sequence YINQMPQKSRE (MW: 1,393.774 Da) and the other peptide showed amino acid sequence YINQMPQKSREA (MW: 1,464.813). Despite this study not was evaluated the antimicrobial activity, but was demonstrated the antioxidant activity, ACE inhibitory and/or antidiabetic activities these peptides.

The physicochemical parameters evaluated showed which the amino acid sequence of Ovipin and molecular weight, YVSPVAIVKGLNIPL and 1581.9494 Da, demonstrated this peptide has one positive charge, your grand average hydropathicity (GRAVY) is 1,193 positive value suggested this peptide be hydrophobic and it showed hydrophobic ratio of 53%. The study Pimchan *et al*., (2023), was identified three peptides with 41, 55 and 62% of hydrophobicity, respectively. Net positive charge and hydrophobicity are important factors and influence the antimicrobial activity because there initial electrostatic between positive charges of the AMPs and negative charges of cell membranes of microorganisms, and than hydrophobic side chains binds to the hydrophobic core, causing cell lysis following membrane disruption and death of microorganism (Pimchan *et al*., 2023; Kopiasz *et al*., 2021). The Ovipin peptide has a positive charge and hydrophobic nature this characteristic to contribute for increase the interaction and facilitate its insertion into the microorganisms membrane, thus causing antimicrobial effects (Schmidtchen, Pasupuleti, Malmsten, 2014).

So this study report Ovipin peptide isolated from chicken eggs *Gallus gallus*, it active against Gram-positive bacteria, Filamentous fugi and Yeast, and amino acid seguence charcterized. Chicken eggs are widely studied and there are several studies that have identified compounds with different functions, mainly with antimicrobial action to different microorganisms. But not have been explored compounds in chicken eggs with activity against *Cryptococcus neoformans* strains, as presented in the present work demonstrating the relevance of the Ovipin peptide for conventional therapies against this microorganism. The next steps is broaden antimicrobial activity of the Ovipin peptide with others microorganisms, such as bacteria, fungi, viruses and parasites, as well as evaluating the secondary structure and the mode of action this peptide.

## Conclusions

Whereas antimicrobial resistance has become a serious concern public health issue because it limits treatment options, the identification and characterization of new antibiotics agents can lead to the development of new compounds that act against resistant pathogenic microorganisms. In this work we reported a novel antimicrobial peptide that was extracted from chicken eggs *Gallus gallus domesticus*, purified by RP-HPLC and characterized by amino acid sequencing. The Ovipin peptide was active against *Micrococcus luteus, Aspergillus niger* and *Cryptococcus neoformans*, an important yeast for medicine because act mainly imunossupressed individuals. Further studies aimed at evaluating the activity of the Ovipin peptide against others multidrug-resistance bacteria, fungi, viruses, and parasites are needed, as well as evaluating secondary structure and the mechanism of action against *C. neoformans*, to understand in which possible virulence factor of this yeast the Ovipin peptide may be acting and that inhibit its growth, and it can be used as a future candidate in the biotechnology or pharmaceutical industry.

## Supporting information

Sup_met

## Author Contributions

Conceptualization, SRS, AM and PISJ; Data curation, SRS; Formal analysis, AM and PISJ; Funding acquisition, AM and PISJ; Investigation, SRS; Methodology, AM and PISJ.; Project administration, AM and PISJ; Resources, PISJ; Supervision, PISJ and AM; Validation, SRS, AM and PISJ; Writing-original draft, SRS; Writing-review and editing, SRS, AM and PISJ.

## Ethical Approval

This research was approved and performed in accordance with the Ethical Principles in Animal Research adopted by the Ethics Committee for the Use of Animals of UNIFESP - CEUA No 9183300118 and Plataforma Brasil CAAE N° 18794419.7.0000.0065 and was approved.

## Data Availability Statement

The data that support the findings of this study are presented in the main manuscript or in the supplementary material of this article.

## Acknowledgments

We would like to thank Caroline Correa, Marta Gomes, Rogerio Lauria and Sinval Gregorio laboratory technicians from the Biophysics Department at Unifesp for all their help in the laboratory. Professor Dr. Edgar Julian Paredes Gamero, and his students Heron and Wagner at Unifesp, for the opportunity to learn about the development of the antitumor study. *Cryptococcus neoformans* VNI (WM148) was kindly provided by Dra Márcia Melhem - Instituto Adolfo Lutz - São Paulo/SP, Brasil.

